# Major Metabolites from *Hypericum Perforatum* L., Hyperforin and Hypericin, are both active against Human Coronaviruses

**DOI:** 10.1101/2024.04.09.588755

**Authors:** I. Raczkiewicz, C. Rivière, P. Bouquet, L. Desmarets, A. Tarricone, C. Camuzet, N. François, G. Lefèvre, J. Samaillie, F. Silva Angulo, C. Robil, F. Trottein, S. Sahpaz, J. Dubuisson, S. Belouzard, A. Goffard, K. Séron

**Affiliations:** Univ. Lille, CNRS, Inserm, CHU Lille, Institut Pasteur de Lille, U1019 – UMR9017 – Center for Infection and Immunity of Lille (CIIL), F-59000 Lille, France; BioEcoAgro, Joint Research Unit 1158, Univ. Lille, INRAE, Univ. Liège, UPJV, YNCREA, Univ. Artois, Univ. Littoral Côte d’Opale, ICV – Institut Charles Viollette, F-59650 Villeneuve d’Ascq, France

## Abstract

COVID-19 pandemic has highlighted the need of antiviral molecules against coronaviruses. Plants are an endless source of active compounds. In the current study, we investigated the potential antiviral effects of *Hypericum perforatum* L.. Its extract contained two major metabolites belonging to distinct chemical classes, hypericin (HC) and hyperforin (HF). First, we demonstrated that HC inhibited HCoV-229E at the entry step by directly targeting the viral particle in a light-dependent manner. While antiviral properties have already been described for HC, the study here showed for the first time that HF has pan-coronavirus antiviral capacity. Indeed, HF was highly active against Alphacoronavirus HCoV-229E (IC_50_ value of 1.10 µM), and Betacoronaviruses SARS-CoV-2 (IC_50_ value of of 0.24 to 0.98 µM), SARS-CoV (IC_50_ value of 1.01 µM) and MERS-CoV (IC_50_ value of 2.55 µM). Unlike HC, HF was active at a post-entry step, most likely the replication step. Antiviral activity of HF on HCoV-229E and SARS-CoV-2 was confirmed in primary human respiratory epithelial cells. Furthermore, *in vitro* combination assay of HF with remdesivir showed that their association was additive, which was encouraging for a potential therapeutical association. As HF was active on both Alpha- and Betacoronaviruses, a cellular target was hypothesized. Heme oxygenase 1 (HO-1) pathway, a potential target of HF, has been investigated but the results showed that HF antiviral activity against HCoV-229E was not dependent on HO-1. Collectively, HF is a promising antiviral candidate in view of our results and pharmacokinetics studies already published in animal models or in human.

## INTRODUCTION

The recent COVID-19 pandemic caused by severe acute respiratory syndrome coronavirus 2 (SARS-CoV-2) has highlighted the urge for broad-spectrum antivirals. Before the emergence of SARS-CoV-2, no specific coronavirus (CoV) antiviral agent was available. Four years after the beginning of the outbreak, we are still lacking for therapeutical options. To date, there are only three direct-acting antivirals (DAA) approved by FDA (Food and Drug Administration) for clinical usage. The ritonavir-boosted nirmatrelvir (Paxlovid®) is the first line-therapy for patients with high risk of developing severe COVID-19. Nirmatrelvir (PF-07321332) is an oral protease inhibitor that is active against the main protease (M^PRO^) of SARS-CoV-2 (1). Ritonavir serves as a booster, since it increases the plasma concentration of nirmatrelvir by inhibiting the cytochrome P450 3A4 (2). The second-line therapy, remdesivir is often given when there is contraindication to Paxlovid®. Remdesivir (GS-5734), a viral RNA-dependent RNA polymerase (RdRp) inhibitor initially developed to treat Ebola infections, was one of the first molecule to demonstrate an antiviral activity against CoV *in vitro* and *in vivo* (3–5). Given intravenously, remdesivir which is an adenosine nucleotide analog prodrug, acts as a chain-terminator (6). The third-line therapy, molnupiravir (MK-4482 or EIDD-2801), a RdRp inhibitor originally developed against hepatitis C virus (HCV) infection, is an orally available prodrug of the ribonucleoside analog EIDD-1931 (β-D-N4-hydroxycytidine) that has a broad-spectrum antiviral activity against RNA viruses (7, 8). Even though these therapies are widely used, some concerns remain. Firstly, even if they are still active against the current circulating variants, the increased use of antivirals supports the development of drug resistance (9, 10). One of the main strategies to avoid the emergence of resistant mutants is the use of combination therapies that allow complete viral clearance, especially in immunocompromised patient (11–13). Secondly, 2% of Paxlovid® treated patients have exhibited a viral rebound, which is not significantly different from the placebo group (14, 15). Thirdly, the poor bioavailability of remdesivir when taken orally makes it challenging for ambulatory care and limits its usage for hospital settings. Fourthly, although molnupiravir is still approved by FDA, it has been withdrawn by the European Medicines Agency (EMA) because the benefice-risk balance was unfavorable for the patient (16). Finally, molnupiravir could pose a risk to the host, as it has been shown to have mutagenic potential in human cells (17). In light of these challenges, the development of new antivirals is more than necessary.

CoVs are positive single stranded-RNA viruses belonging to the *Coronaviridae* family, the *Orthocoronavirinae* subfamily and the Nidovirales order (18). Seven CoV infect human (HCoV) so far and can be distinguished into two groups based on the clinical presentation, from mild, HCoV-229E, -OC43, -HKU1, and -NL63, to severe symptoms, SARS-CoV, Middle-East respiratory syndrome coronavirus (MERS-CoV) and SARS-CoV-2 (19). Coronaviruses are enveloped RNA viruses. Structural proteins spike (S), envelope (E), and membrane (M) proteins are embedded into the envelope lipid bilayer and protect the viral genome which is associated with nucleoprotein (N). The S protein mediates the host-cell attachment and the viral entry by recognizing host-specific receptors (20). The virus enters into the cell via two different pathways, depending on the expression of cellular proteases on the cell surface, such as TMPRSS2 (21). If the latter is expressed, the S protein is cleaved at the cell surface and the viral particle fuses with host plasma membrane. The endosomal pathway is the second entry pathway and the fusion occurs with the endosomal membrane (22). The genome is then released into the cytosol, where it is translated into two polyproteins, pp1a and pp1ab. These two are cleaved by the papain-like protease nsp3 (PL^pro^) and the main protease nsp5 (M^PRO^), into several nonstructural proteins (nsp) such as nsp12, the viral RdRp, engaging the replication step (23). Then the virus is assembled and secreted.

According to World Health Organization, around 80% of the population relies on herbal medicine to heal themselves, and plants are a great source of active molecules with huge structural diversity. Several natural products have been shown to exhibit antiviral activity against viruses of different families *in vitro* (24). We have recently shown that pheophorbide a (Pba), a chlorophyll degradation product is active against SARS-CoV-2 and MERS-CoV (25), that the red algae derivate griffithsin is active against MERS-CoV (26), and that some cinnamoyl oleanolic acids isolated from *Hippophae rhmnoides* are active against both SARS-CoV-2 and HCoV-229E (27). Also, we believe that plants could contain promising antiviral molecules of different chemical classes that are still unknown.

*Hypericum* Tourn ex L. is a cosmopolitan genus with 512 recognized species to date (28). *Hypericum perforatum* L. (perforated Saint John’s wort, SJW), belonging to Hypericaceae, is the most common and well-known species among this genus (29). Its chemical composition includes flavonols (quercetin and its glycosides including hyperoside, rutin, isoquercitrin, quercitrin, miquelianin), biflavonoids (I3,II8-biapigenin and amentoflavone), naphtodianthrones (hypericin (HC) and its analogues), prenylated phloroglucinols (hyperforin (HF) and its analogues) (30). Some standardized extracts of SJW, have been studied in many clinical trials to assess their effectiveness in depression also demonstrating a good tolerability (31–33). However, HC appears to induce adverse effects such as phototoxicity (34) and HF is known to be an enzyme inducer (31). HF and HC are the active compounds of *Hypericum perforatum* L. known to be responsible for its anti-depressive properties. To date, only HC has been described for antiviral activity against enveloped viruses such as human immunodeficiency virus 1, Sindbis virus and murine cytomegalovirus with a light-dependent activity (35). More recently, an antiviral activity against SARS-CoV-2 has been discovered for this naphtodianthrone (36).

Here we highlighted that two major metabolites of *Hypericum perforatum* L., HC and HF, exhibit antiviral activity against HCoV, with distinct modes of action with HC inhibiting entry of HCoV an HF a post-entry step. Furthermore, our data described for the first time an antiviral capacity for HF with pan-coronavirus activity.

## RESULTS

### 1. Hypericum perforatum *L.* and its metabolites, HF and HC, are active against HCoV-229E

In order to identify new antiviral compounds against HCoVs, we have selected the crude methanolic extract of *Hypericum perforatum* L. based on a previous screening of several plants. Its antiviral activity was tested against HCoV-229E-Luc in a dose-response assay. The results showed that the crude methanolic extract inhibited HCoV-229-Luc infection in a dose-dependent manner with 50% inhibitory concentration (IC_50_) of 18.73 ± 0.84 µg/mL (**Figure 1A**). No cytotoxicity was observed at active concentrations (**Figure 1A**). The results suggested the presence of at least one active compound within *Hypericum perforatum* L. crude extract which is known to contain two major metabolites, HF, a prenylated phloroglucinol derivative, and HC, an anthraquinone derivative. To determine if these compounds were responsible for the anti-coronavirus activity, dose-response and cytotoxicity assays were performed. The results showed that the two metabolites exhibited antiviral activity against HCoV-229-Luc with IC_50_ values of 0.37 ± 0.02 µM and 1.10 µM ± 0.08 µM, respectively (**Figures 1B and C**). Cytotoxicity tests showed that both compounds are not toxic at active concentrations with a 50% cytotoxic concentration (CC_50_) values of 25.77 ± 2.58 µM and 19.35 ± 2.08 µM for HC and HF, respectively, resulting in a selective index (SI) of 69 for HC and 17 for HF (**Figures 1B and C**).

**Figure 1.**
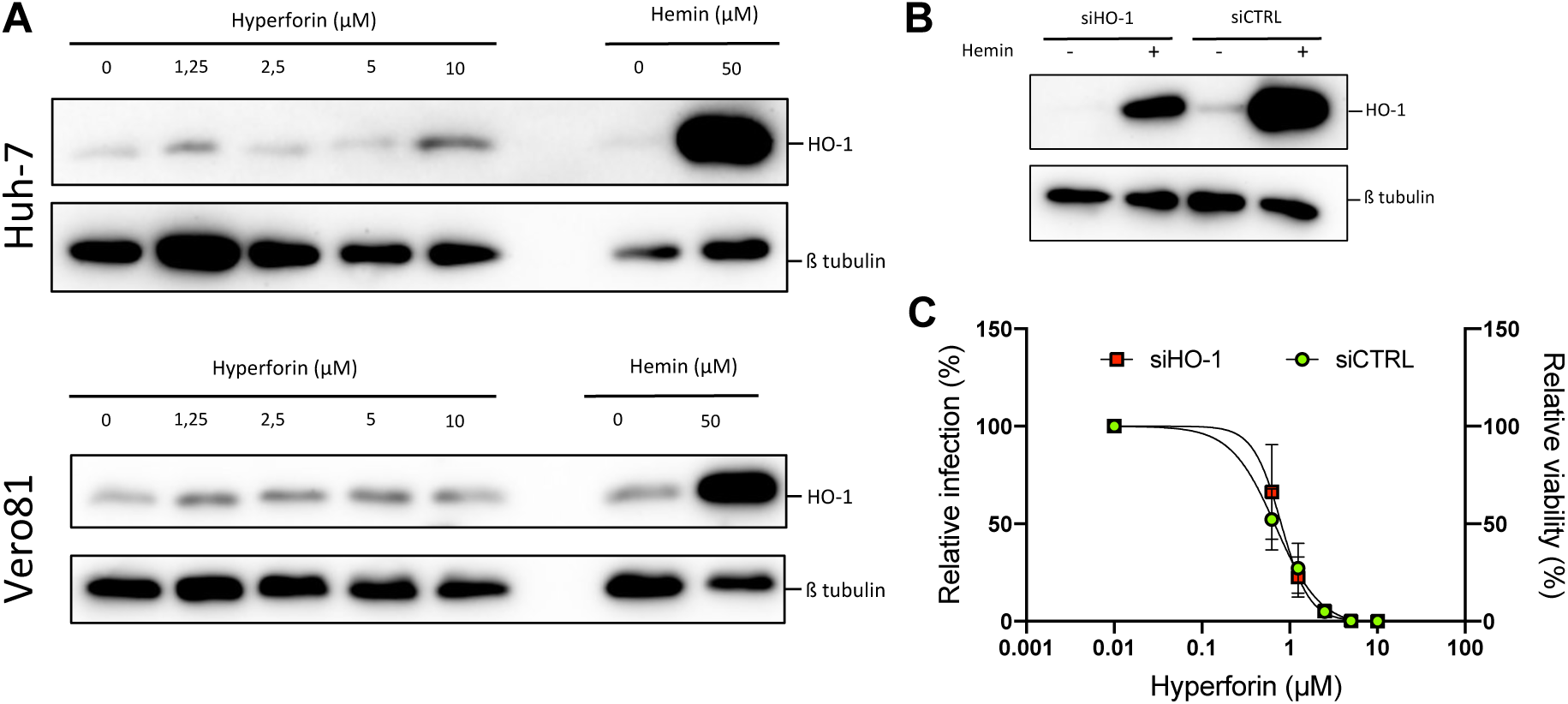
The antiviral activity of HF is not linked with HO-1 pathway. **A)** Huh-7 cells or Vero81 cells were treated with increasing concentrations of HF. HO-1 expression was then studied by Western Blot. **B)** Huh-7 cells were treated with siRNA targeting HO-1 (siHO-1) or control siRNA (siCTRL). 24 h later, cells were treated or not with 50 µM of hemin. HO-1 expression was then studied by Western Blot. **C)** Dose-response assays of HF against HCoV-229E were performed in presence of siHO-1 or siCTRL. The luciferase activity was measured after 24 h of incubation. Western Blots were representative for 3 independent experiments. Data were presented as means +/- SEM of 3 independent experiments performed in triplicates.

### 2. HC inhibits the entry of HCoV-229E in a light-dependent manner and by directly targeting the viral particle

We further investigated each active compound individually to determine their mechanism of action. It was recently underlined that HC inhibited SARS-CoV-2 at the entry step (36). To characterize the mechanism of action of HC on HCoV-229E, a time-of-addition assay was performed by adding the compound at 4 µM (corresponding to the IC_90_) at different timepoints during infection, either before, during or after the inoculation of the cells by the virus (**Figure 2A**). The results showed that HC significantly reduced HCoV-229E-Luc infection when added from pre-treatment condition to 2 h post-inoculation (p.i.), meaning that it inhibited the infection at an early step (**Figure 2B**), probably the entry step. An entry assay was performed with particles pseudotyped with HCoV-229E S protein (229pp) and mimicking virus entry, in the presence of HC. The results showed that HC significantly decreased 229Epp entry in a dose-dependent manner (**Figure 2C**). Taken together, these results highlighted that HC is an inhibitor of HCoV-229E entry, which is consistent with already published data on SARS-CoV-2.

**Figure 2.**
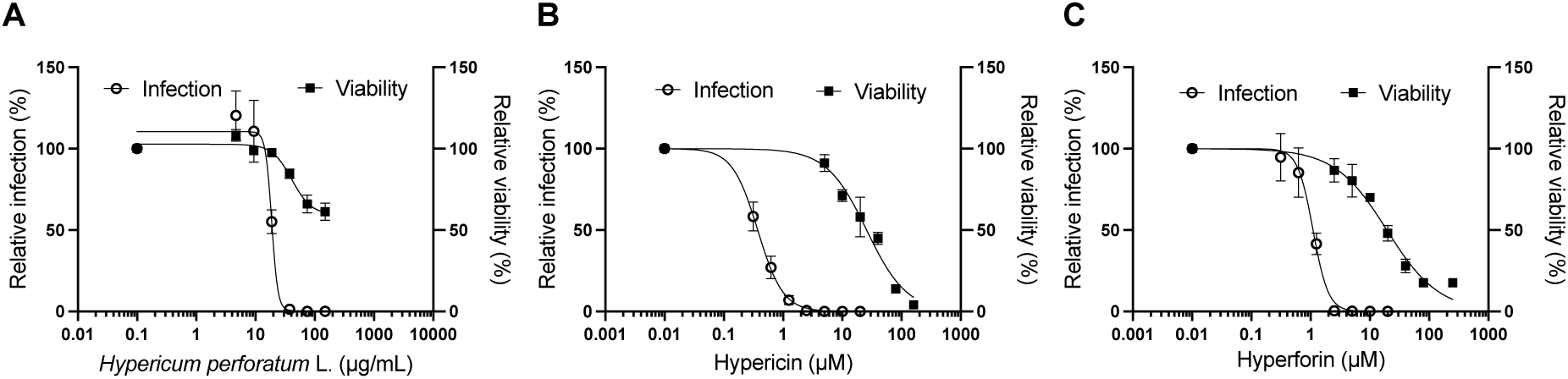
Cytotoxicity and antiviral activity of *Hypericum perforatum* L. and its metabolites against HCoV-229E. Huh-7 cells were inoculated with HCoV-229E-Luc in the presence of increasing concentrations of (**A**) *Hypericum perforatum* L. crude methanolic extract, (**B**) HC or (**C**) HF. Cells were lysed 7 h after infection and luciferase activity was quantified. For the cytotoxicity assay, cells were incubated 24 h with increasing concentrations of the crude extract or the molecules and MTS assay were performed. Results are expressed as the means +/- SEM of 3 independent experiments performed in triplicate.

To inhibit viral entry, a compound may act directly on the viral particle before its attachment to the cell surface. To assess if HC was such inhibitor, we incubated HCoV-229E-Luc with HC at high concentration (2 µM) for 30 min. Then, before inoculation, we diluted the mixture 10 times to reach a final concentration of HC of 0.2 µM. This concentration is inactive on HCoV-229E. As controls, cells were inoculated by HCoV-229E-Luc in the presence of HC either at “high” concentration (2 µM) and “low” concentration (0.2 µM). It is important to note that the MOI was kept constant in all conditions. The results showed no significant inhibition of infection in presence of 0.2 µM HC, whereas a similar significant inhibition of infection was observed for the condition “2 µM > 0.2 µM” HC and infection with 2 µM HC (**Figure 2D**). Taken together these results showed that HC targeted the viral particle before infection. It is well known that HC is a photoactivable molecule and has a light-dependent activity on several viruses (35). To determine if antiviral activity of HC on HCoV-229E is light-dependent, a dose-response assay was carried out in the presence or absence of light. The white visible light is sufficient to photoactivate HC, so we used the light of the safety cabinet (37). The results confirmed the dose-dependent inhibition of infection of HCoV-229E-Luc in the presence of HC under normal light condition (**Figure 2E**) similar to the results presented in **Figure 1B**. However, when the experiment was carried out in the dark, no inhibition of HCoV-229E-Luc infection was observed even at high HC concentration up to 5 µM (a concentration above IC_90_ under light exposure) (**Figure 2E**). These results clearly demonstrated that HC directly targets the viral particle and inhibits HCoV-229E entry in a light-dependent manner.

### 3. HF has a pan-coronavirus antiviral activity

HF is less studied than HC in literature and no data are available on its potential antiviral activity. Thus, we performed antiviral assays against three highly pathogenic HCoVs, SARS-CoV, MERS-CoV and SARS-CoV-2. For SARS-CoV-2, we had access to different variants, the initial Wuhan strain (D614), alpha variant B1.1.7 and omicron variant B1.1.529. Dose-response inhibition studies were conducted and the results were shown in **Figure 3**. Antiviral assays on SARS-CoV-2 variants were performed in Vero-81/TMPRSS2 cells and antiviral activity was assessed by viral titration. The results showed that HF inhibited infection of all tested SARS-CoV-2 variants with calculated IC_50_ values of 0.98 ± 0.28 µM, 0.24 ± 0.02 µM, and 0.29 ± 0.13 µM for strain D614, alpha variant B1.1.7 and omicron variant B1.1.529 respectively, without any cytotoxicity at the active concentration (CC_50_ = 45.91 µM) (**Figure 3A**). The respective SI were all higher than 40 (**Table 2**). These data were confirmed by Western blot analyses in human lung A549/ACE2 cells infected with SARS-CoV-2 alpha variant (**Figure 3B**). A dose-dependent decrease of the expression of SARS-CoV-2 N protein was observed in presence of HF. Dose-response assays were also conducted against the two others highly pathogenic HCoVs, SARS-CoV and MERS-CoV, in Vero-81/TMPRSS2 and Calu-3 cells, respectively. Our results highlighted that HF is highly active against both SARS-CoV and MERS-CoV with an IC_50_ values of 1.01 ± 0.12 µM and 2.55 ± 0.28 µM, respectively, resulting in SI of 45 for SARS-CoV (**Figure 3C and 3D**, **Table 2**). CC_50_ value in Calu-3 cells was not determined precisely, but, as observed in **Figure 3D**, it was higher than 20 µM, with SI>7. Taken together, these data suggested that HF has a pan-coronavirus antiviral activity with IC_50_ values ranging from 0.24 to 2.55 µM.

**Figure 3.**
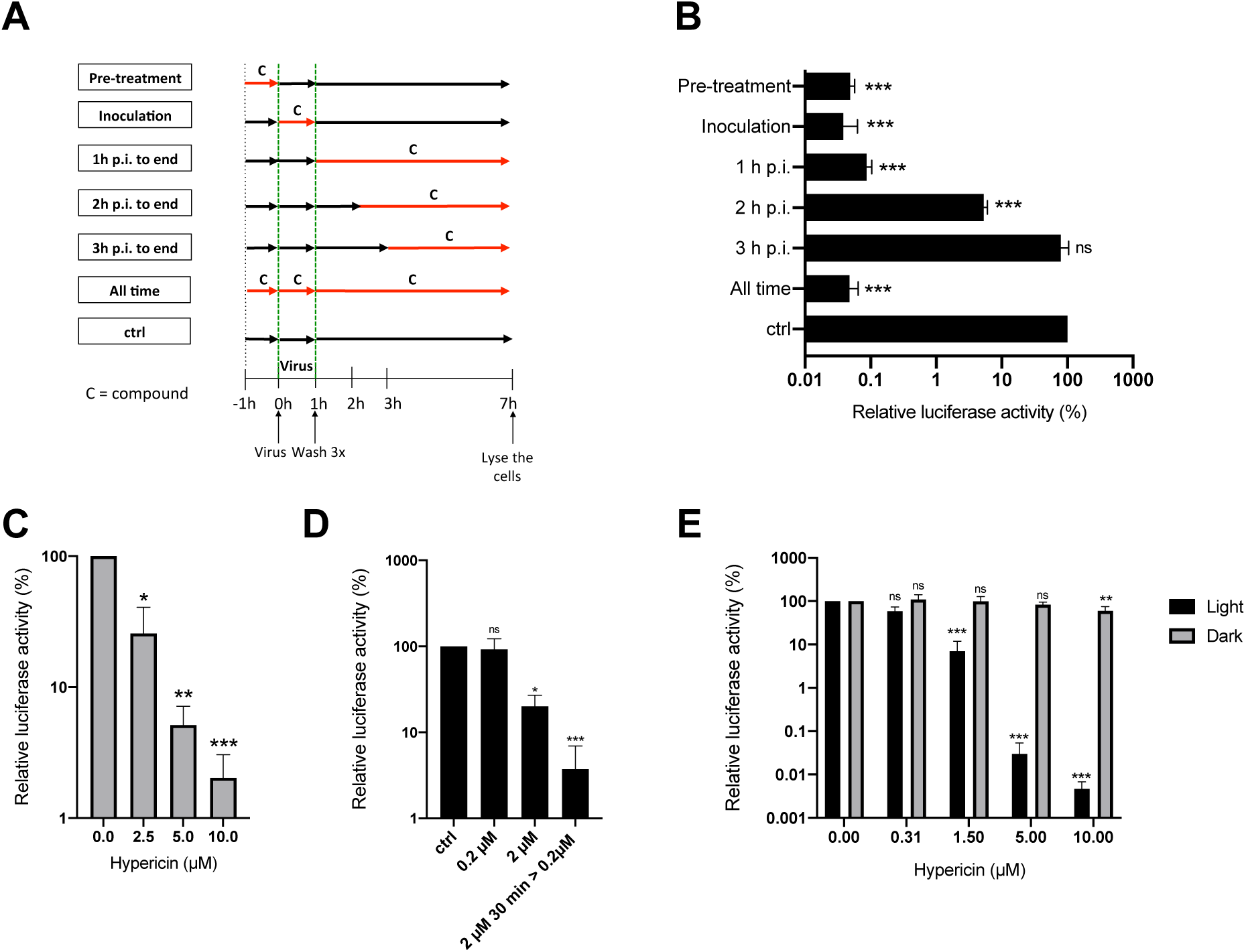
HC inhibits HCoV-229E entry by directly targeting the virus in a light-dependent manner. **A)** Graphical representation of the time-of-addition assay. The compound (C) was added at different time points during infection. **B)** Huh-7/TMPRSS2 cells were inoculated with HCoV-229E in the presence of 4 µM HC at different timepoints. Cells were lysed 7 h p.i. and luciferase activity was quantified. **C)** Huh-7/TMPRSS2 cells were inoculated with 229Epp in the presence of increasing concentrations of HC. After 2 h, the inoculum was removed and replaced with medium without compound for 46 h. Cells were lysed and luciferase activity was quantified. **D)** HCoV-229E-Luc was incubated with 2 µM HC for 30 min and dilute 10 times to reach a concentration of 0.2 µM HC for inoculation of Huh-7/TMPRSS2 cells (2 µM > 0.2 µM). As controls, cells were inoculated with HCoV-229E-Luc in the presence of 0.2 or 2 µM HC. The MOI was the same in all conditions. Cells were lysed 7 h p.i. and luciferase activity was quantified. **E)** Huh-7/TMPRSS2 cells were treated with increasing concentrations of HC and inoculated with HCoV-229E-Luc in dark or light condition. Plates were kept in the dark or exposed to the light for 10 min during inoculation, and 10 min p.i.. Cells were lysed as described and luciferase activity was quantified. Results were expressed as the means +/- SEM of 3 independent experiments performed in triplicate. Statistical analyses were performed using Mann-Whitney non-parametric test; ns: non significative, * *P* < 0.05, ** *P* < 0.01, *** *P* < 0.001. C: compound. p.i.: post inoculation.

**Table 1.** Cytotoxicity and antiviral activity of HF against HCoVs. ND: not determined.

| Virus | Cells | IC<sub>50</sub> (μM) | CC<sub>50</sub> (μM) | SI |
| --- | --- | --- | --- | --- |
| HCoV-229E | Huh-7/TMPRSS2 | 1.10 ± 0.08 | 19.35 ± 2.08 | 17 |
| SARS-CoV-2 (D614) | Vero-81/TMPRSS2 | 0.98 ± 0.28 | 45.91 ± 4.85 | 46 |
| SARS-CoV-2 alpha (B1.1.7) | Vero-81/TMPRSS2 | 0.24 ± 0.02 | 45.91 ± 4.85 | 188 |
| SARS-CoV-2 omicron (B1.1.529) | Vero-81/TMPRSS2 | 0.29 ± 0.13 | 45.91 ± 4.85 | 158 |
| SARS-CoV | Vero-81/TMPRSS2 | 1.01 ± 0.12 | 45.91 ± 4.85 | 45 |
| MERS-CoV | Calu-3 | 2.55 ± 0.28 | > 20.0 | ND |

**Table 2.** Synergy scoring obtained from Synergy Finder for HF and remdesivir combination.

| Model | Synergy score | *p*-value |
| --- | --- | --- |
| ZIP | 4.23 | 4.44e-04 |
| Loewe | 3.09 | 5.22e-03 |
| HAS | 7.05 | 5.42e-23 |
| Bliss | 5.13 | 9.11e-07 |

### 4. HF is an inhibitor of HCoV replication step

To characterize the mechanism of action of HF against HCoVs, time-of-addition assays against HCoV-229E and SARS-CoV-2 were performed. The data presented in **Figure 4A** and **4B** showed a higher inhibition of infection when HF was added at 1 h to 3 h p.i., for both HCoV-229E-Luc and SARS-CoV-2. Although HF significatively inhibited the infection of HCoV-229E at all steps (from pre-treatment to 3 h p.i.), it decreased the infection by more than 2xLog_10_ from 1 h p.i. to 3 h p.i. (**Figure 4A**). Similar results were observed with SARS-CoV-2; N protein was not detected when HF was added from 1 h to 3 h p.i. (**Figure 4B**). Pba, a natural compound targeting the viral envelope, and GC376, a protease inhibitor, were added as an entry and a replication inhibitor, respectively. Similar profiles of N expression were observed with GC376 and HF. Chloroquine was used to control the expression of TMPRSS2. These data suggested that HF is an inhibitor of the replication step of both SARS-CoV-2 and HCoV-229E.

**Figure 4.**
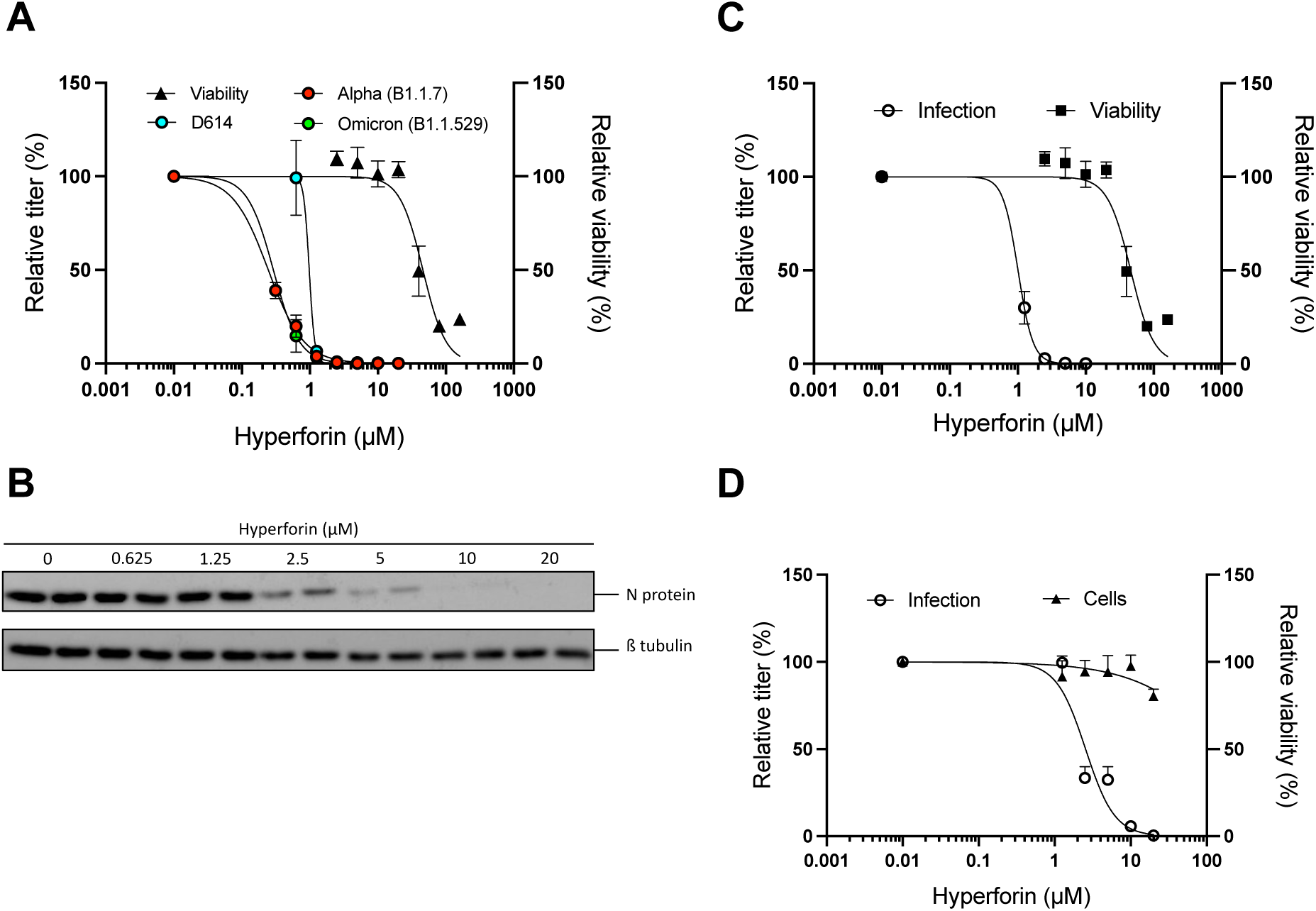
Antiviral activity of HF against several HCoVs. **A)** Vero-81/TMPRSS2 cells were inoculated with different variants of SARS-CoV-2 (D614 strain, alpha or omicron variant) in the presence of increasing concentrations of HF for 16 h. Supernatants were collected, and the viral titers were determined by TCID50/mL. Viability at 24 h was assessed by MTS assay. **B)** A549/ACE2 cells were infected with SARS-CoV-2 alpha variant in the presence of increasing concentrations of HF for 16 h. Cells were lysed and Western Blot analysis was performed. Western blot is representative of 3 independent experiments. **C)** Vero-81/TMPRSS2 cells were inoculated with SARS-CoV in the presence of increasing concentrations of HF for 16 h. Supernatants were collected for viral titration. Viability was determined by MTS assay. **D)** Calu-3 cells were inoculated with MERS-CoV in the presence of increasing concentrations of HF for 16 h. Supernatants were collected for viral titration; Cell number was determined by staining of the nuclei with DAPI. Data were presented as means +/- SEM of 3 independent experiments performed in duplicate (**A, C, D**).

To confirm this hypothesis, entry inhibition assays were performed with either 229Epp and SARS2pp. No significant decrease of infection was observed for any of the particles in the presence of HF up to 5 µM (**Figure 4C**) confirming that HF is not an entry inhibitor. Altogether these data suggested that HF is an inhibitor of HCoV post-entry step, most likely the replication step.

### 5. HF inhibits HCoV-229E and SARS-CoV-2 infection in human primary epithelial cells

To gain relevance, the antiviral activity of HF was then tested in human primary airway cells cultivated at air-liquid interface, considered as a preclinical model for human coronaviruses. First, the cytotoxicity of HF in human airway epithelia (MucilAir^TM^-HAE) was determined by measuring LDH secretion and TEER at 24 and 48 h. LDH secretion higher than 5 % and TEER lower than 100 Ω.cm^2^ reflect damaged cells. No cytotoxicity was observed with 4 µM HF at 24 and 48 h for the two measured parameters (**Figures 5A and B**). However, cytotoxicity was observed with 12 µM HF at 48 h, with LDH secretion higher than 5% compared to control and TEER lower than 100 Ω.cm^2^ (**Figures 5A and B**). Thus, for antiviral assays, two HF concentrations, 4 and 12 µM, were tested against HCoV-229E-Luc and only one, 4 µM, against SARS-CoV-2 due to a longer incubation time of 48 h.

**Figure 5.**
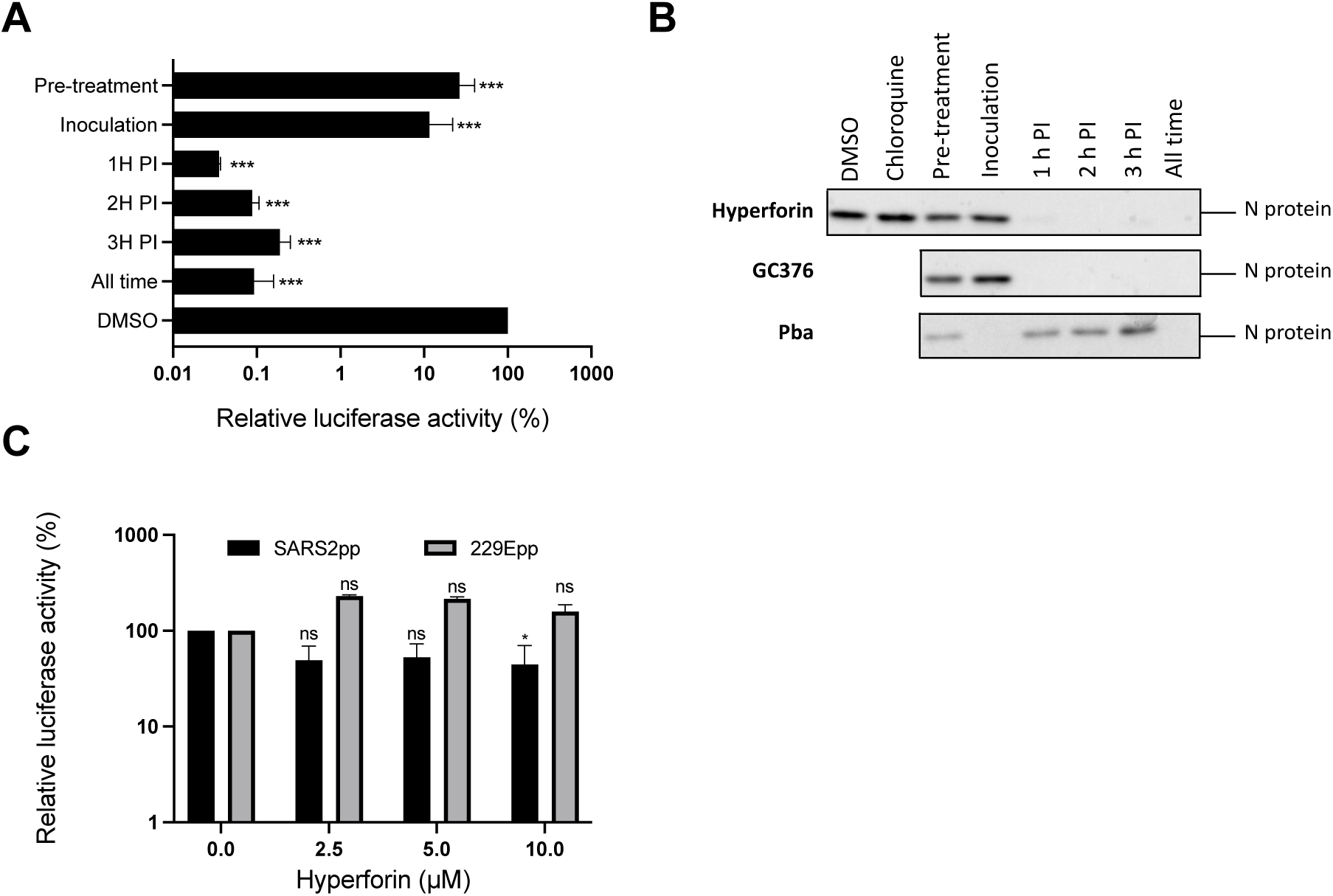
HF inhibits a post-entry step of the viral cycle. **A)** Time-of-addition assay was performed against HCoV-229E in Huh-7/TMPRSS2 cells in the presence of 4 µM HF as previously described in Figure 2A. **B)** Time-of-addition assay was performed against SARS-CoV-2 in Vero-81/TMPRSS2 cells in the presence of 20 µM HF, or 10 µM GC367 and 1 µM Pba, as controls. 10 µM chloroquine was used as a control of TMPRSS2 expression. Cells were lysed 16 h post-infection and lysates subjected to Western blot analysis as previously described. Western blot is representative for 3 independent experiments. **C)** HEK-293TT/ACE2 and Huh-7/TMPRSS2 cells were inoculated with SARSpp and 229Epp, respectively in the presence of increasing concentrations of HF for 2 h. Inoculum was removed and replaced with medium without compound for 46 h. Cells were lysed and luciferase activity was quantified. Data were expressed as mean +/- SEM of 3 independent experiments performed in triplicate. Statistical analyses were performed using the Mann-Whitney nonparametric test. n.s. not significant. * *P* < 0.05, *** *P* < 0.001.

HAE were inoculated with HCoV-229E-Luc in the presence of 4 or 12 µM HF and 10 µM GC376 as a control, for 24 h. Viral RNA at the apical surface was quantified as well as luciferase activity in cell lysates. The results showed a decrease in viral RNA copies and luciferase activity when cells were treated with 12 µM HF (**Figures 5C and D**), similar to the decrease observed with 10 µM GC376. These results highlighted that HF is active against HCoV-229E in human primary cells.

To determine if HF could also inhibit SARS-CoV-2 in this preclinical model, HAE were inoculated with the virus in the presence of 4 µM HF. Viral infectious titers at the apical surface and intracellular viral RNA were quantified (**Figures 5E and F**). The data showed a decrease of both viral RNA copies and infectious titers in the presence 4 µM HF. A 3xLog_10_ decrease in viral titers was observed upon 4 µM HF treatment similar to the decrease observed with 5 µM remdesivir demonstrating that HF is an inhibitor of SARS-CoV-2 infection in HAE.

Taken together, these data underlined that HF is active against HCoVs in human primary respiratory epithelial cells grown in air-liquid interface.

### 6. HF has an additive effect with remdesivir

The results presented in this study demonstrate that HF encompasses many characteristics of an antiviral agent to be used in therapy. In order to determine the potential use of HF as therapeutic in clinic, combination studies of HF with remdesivir and nirmatrelvir, were performed. Checkboard assays were performed with double serial dilutions of HF and remdesivir or nirmatrelvir (**Figure 6**).

**Figure 6.**
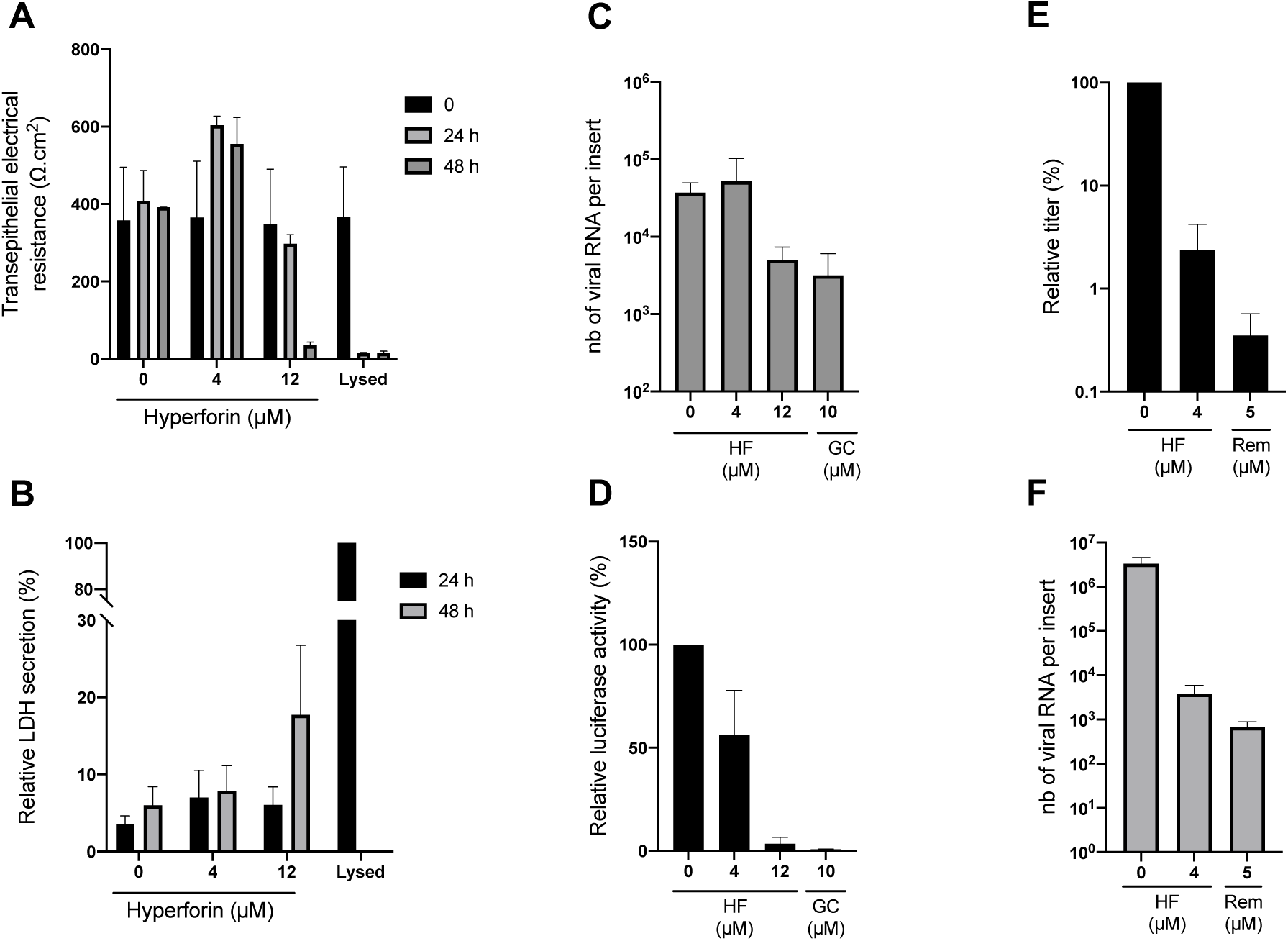
HF is active in human primary respiratory epithelial cells. Cytotoxicity was determined by measuring the TEER (**A**) and LDH secretion (**B**) according to the manufacturer’s recommendations. Cells were inoculated with HCoV-229E-Luc at the apical surface of MucilAir™ HAE in the presence of 4 or 12 µM HF, and 10 µM GC376 (GC) for 24 h. RNA was extracted from apical wash and was quantified by RT-qPCR (**C**). Cells were lysed and luciferase activity was measured (**D**). Cells were inoculated with SARS-CoV-2 at the apical surface in the presence of 4 µM HF or 5 µM remdesivir (Rem) for 48 h. Infectious virus secreted at the apical surface was quantified by TCID50/mL (**E**) and intracellular viral RNA by RT-qPCR analysis (**F**). Data were expressed as mean +/- SEM of 2 independent experiments.

F1G-Red Vero-81 reporter cell line was used to monitor SARS-CoV-2 infection (38). First, the cytotoxicity of each combination was assessed by quantifying the number of nuclei. No cytotoxicity was observed for any of the combinations of HF with remdesivir or nirmatrelvir (**Supplemental Figure 1**). The combination effect was then assessed with SynergyFinderPlus. As shown in **Figure 7C**, the combination of HF with remdesivir is additive, with synergy scores ranging from -10 to +10, with the four mathematical models (HSA, Loewe, Bliss and ZIP) and significant *p*-value (**Table 2**).

**Figure 7:**
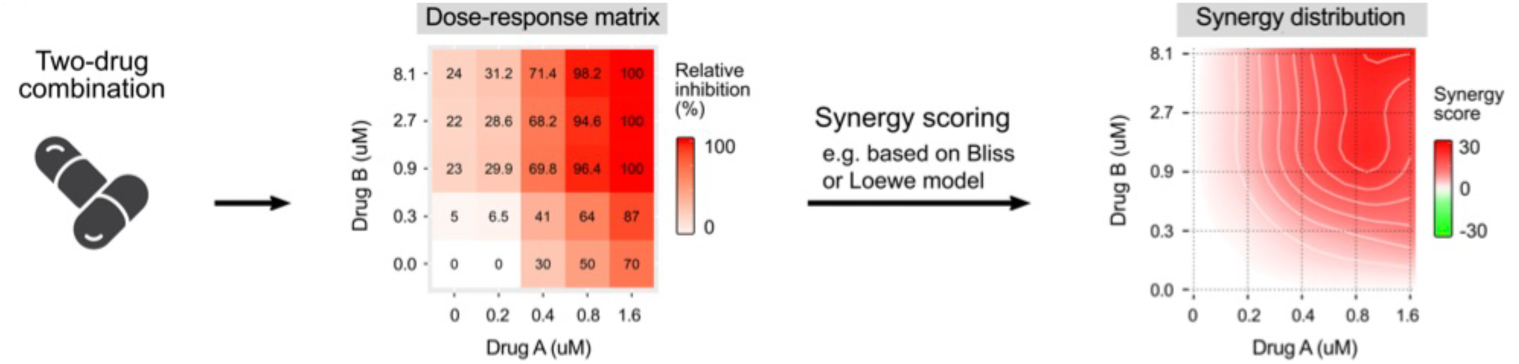
Protocol of the combination assay. Cells (F1G-Red) were treated with increasing concentrations of drug A and drug B and challenged with SARS-CoV-2. The infection was then assessed by confocal microscopy. Nine images per well were taken and infected cell number was quantified. The data were then uploaded on SynergyFinderPlus. Synergy scores were calculated based on 4 mathematical models (HSA, Loewe, Bliss and ZIP). The combination is synergistic if the score is above 10; additive if it is ranging from -10 to 10; and antagonistic if it is below 10.

A synergistic area, score above 10, was highlighted with HF, ranging from 2.5 to 10 µM, combined to remdesivir, ranging from 37.5 to 150 nM, with a synergy score of 14.9 (**Figure 7B**). For nirmatrelvir, none of the mathematical model was able to give a significant synergy score (**Supplemental Table 2**).

### 7. The antiviral activity of HF is not dependent of HO-1 pathway

Due to the pan-coronavirus antiviral activity of HF, we put forward the hypothesis that it may regulate a cellular factor necessary for HCoVs replication. Recently, HF was described as an inducer of heme oxygenase 1 (HO-1) pathway by upregulating the expression of HO-1 (39). Moreover, it was also recently shown that hemin, an HO-1 inducer that upregulates HO-1 expression, has an antiviral activity against SARS-CoV-2 (40). HO-1 pathway was also described to be involved in antiviral immune response against different viruses (41). Thus, we hypothesized that the antiviral activity of HF against HCoV could be linked to the HO-1 pathway. To evaluate this hypothesis, Huh-7 cells or Vero81 cells were treated with increasing concentrations of HF and then HO-1 protein expression was detected by Western Blot. The results showed that HF up to 10 µM (active antiviral concentration; 10 x IC_50_) was not able to upregulate HO-1, unlike hemin, used as a control, which induced a strong up regulation of the protein expression (**Figure 8A**). Next, to further confirm these results, siRNA targeting HO-1 (siHO-1) were used to knock-down its expression prior to infection of Huh-7 cells with HCoV-229E. HO-1 protein expression was strongly decreased by siHO-1, and was weakly induced by hemin (**Figure 8B**). However, similar HF antiviral activities against HCoV-229E were observed in wild-type Huh-7 cells compared to siHO-1 cells (**Figure 8C**). Taken together, the results highlighted that HO-1 pathway is not involved in HF antiviral activity against HCoV-229E.

**Figure 8.**
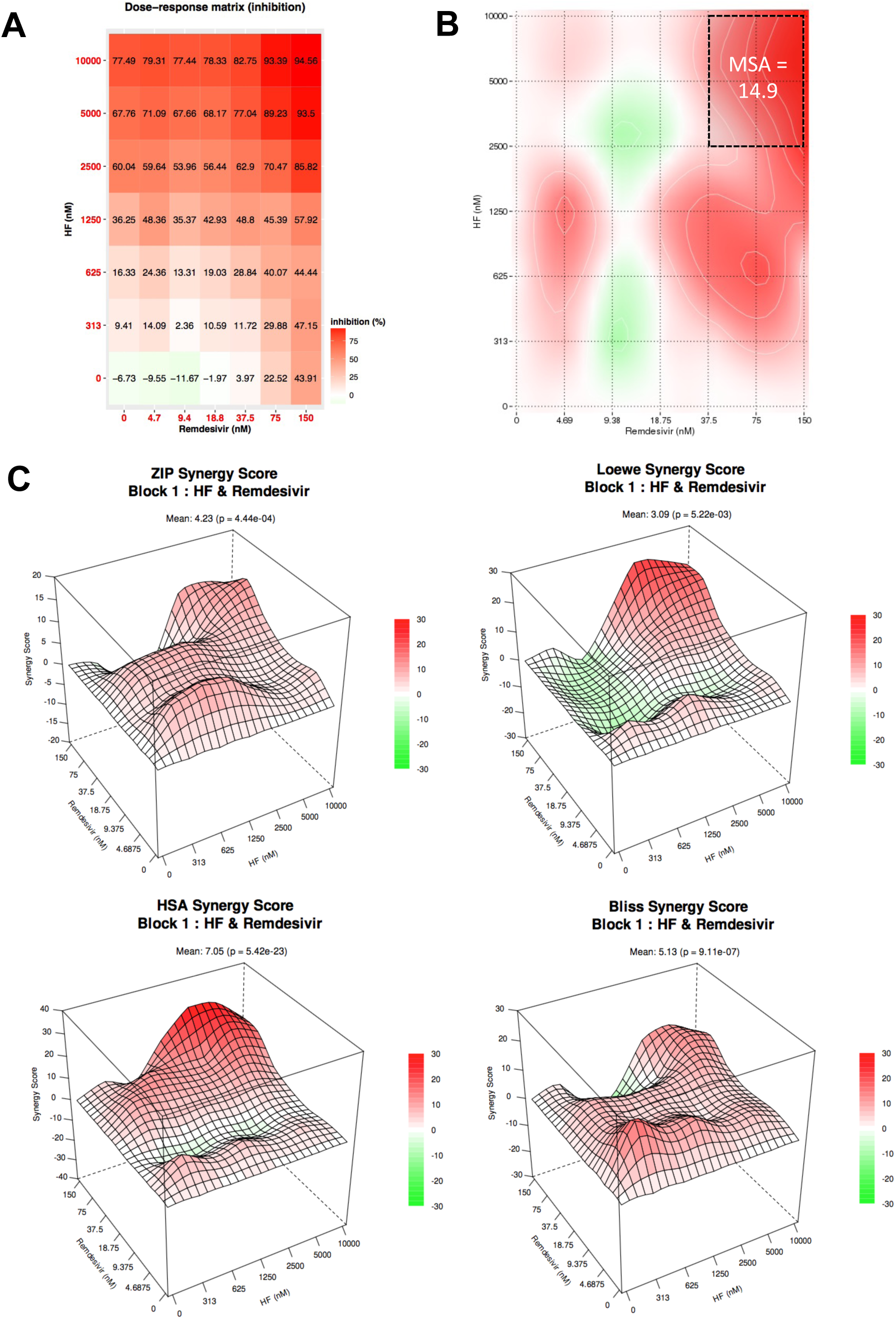
Combination of HF and remdesivir. **A)** Inhibition of infection heatmap. **B)** Most synergistic area (MSA) obtained with SynergyFinder 3.0 for HSA model. **C)** Heatmaps obtained with SynergyFinderPlus.

## DISCUSSION

The development of broad-spectrum antiviral agents has become a necessity to support the pandemic preparedness. Natural compounds are a great source of active compounds with biological activities, including among others anticancer, antibacterial or antiviral properties (42). Although there are no natural antiviral molecules used in clinical settings yet, some plant extracts are currently being tested in clinical trials, like Antiwei granule (a mix of Chinese herbs) for the treatment of influenza (43), *Viola odora* L. against SARS-CoV-2 infection (44) or *Sutherlandia frutescens* (L.) R.Br. against HIV infection (45).

Here, we showed that HF, a major metabolite of SJW (*Hypericum perforatum* L.), a prenylated phloroglucinol, has antiviral capacity against highly pathogenic HCoVs, SARS-CoV, MERS-CoV, and SARS-CoV-2, and the low pathogenic HCoV-229E. Moreover, HF was active in combination with remdesivir and in human primary airway cells. In addition, we highlighted that HF might be an inhibitor of the replication step. Interestingly, HC, the second major metabolite of SJW, a naphtodianthrone, displayed also antiviral capacities against HCoV-229E infection but with a different mechanism of action than HF.

We demonstrated that HC has photo-dependent activity and targeted the viral particle. The light-dependent mechanism of action of HC is consistent with previous reports on other enveloped viruses such as human immunodeficiency virus 1, Sindbis virus and murine cytomegalovirus (35, 46). Nonetheless, its light-dependent mechanism of action and its phototoxicity are not compatible with clinical application in infected patients (34), but can be potentially of interest for an environment disinfectant. Consequently, we focused our study on characterization of HF antiviral activity.

We demonstrated that HF displayed pan-coronavirus activity, a feature that makes this natural compound quite unique in the literature. Without any structural modification, IC_50_ values are close to 1 µM, HCoV-229E (IC_50_ = 1.10 µM), SARS-CoV-2 (IC_50_ = 0.24 – 0.98 µM), SARS-CoV (IC_50_ = 1.01 µM) and MERS-CoV (IC_50_ = 2.5 µM). Pharmacokinetics and bioavailability data are already available in the literature for SJW’s extract. It is important to note that hyperforin is the major metabolite and hypericin is less abundant (up to 4.5% and 0.15% of the extract, respectively) (47). Indeed, different studies show that the circulating concentration of HF is within the range of our IC_50_ values with a maximal concentration of 690 nM and 1100 nM in rats and mice models, respectively, after administration of 300 mg/kg of SJW extract and 5.2 mg/kg of hyperforin, respectively (48, 49). In human, ingestion of a single dose of 300 – 1200 mg SJW’s extract (containing 5 % of HF) led to a detectable concentration of hyperforin in plasma ranging between 200 and 300 nM in healthy volunteers in two different studies (48–50). Moreover, different reports also demonstrated that St John’s Wort extract was neither toxic in animal models nor in humans (31, 47, 51).

Although these results are encouraging, the distribution of HF in lungs after oral administration has not been reported yet. Donà *et al.* (52) showed that HF was able to reduce lung metastases in mice after intraperitoneal injection, demonstrating that the compound was able to reach lungs *in vivo*. Our results in HAE demonstrated that HF is active when administrated at the air interface. It would be interesting to quantify HF in the lungs after intranasal administration. Pharmacokinetics studies are needed to optimize the dose and the route of administration. Moreover, HF or SJW formulation might also be optimized to increase their bioavailability (49, 50).

Besides the two major metabolites, HF and HC, the crude extract of *Hypericum perforatum* L. also exhibited antiviral activity against HCoV-229E and SARS-CoV-2 (our data and (36)). *Hypericum perforatum* L. is one out of more than 500 species of *Hypericum* described so far. The composition in these two metabolites is very variable within these species. It would be very interesting to correlate the antiviral activity of *Hypericum* species with their composition. This will help to determine if the antiviral activity is mainly driven by HF or HC, or if both may have additive or synergistic effect. We showed that HF and HC had different mechanism of action, entry inhibition for HC and translation or replication inhibition for HF. Consequently, it could be expected that these two compounds may have, at least, an additive antiviral effect.

To avoid resistance, it is admitted that antiviral therapy should combine 2 or 3 antiviral agents. Our results showed that HF could be associated with remdesivir, a RdRp inhibitor, with additive antiviral activity, and synergistic at high concentration, *in vitro*. This result is promising, but, unfortunately, no combination of HF with nirmatrelvir, the anti-protease agent of Paxlovid, lacked additive or synergistic effect. However, no antagonist effect was observed, showing that HF could still be envisaged as a promising antiviral agent. More experiments are needed to explain these results.

Here we demonstrated that HF is active against all the highly pathogenic HCoVs described so far, SARS-CoV, MERS-CoV and SARS-CoV-2. They all belong to the Betacoronavirus genus. Interestingly, HF was also active against HCoV-229E which is a member of the Alphacoronavirus genus. It would be interesting to test the antiviral capacity of HF against other human and animal CoVs. HF time-of-addition assay on HCoV-229E and SARS-CoV-2 suggested that HF could inhibit the replication step. Indeed, HF has a similar kinetic profile as GC-376, a SARS-CoV-2 Mpro inhibitor. We thus hypothesized that HF might target a cellular factor necessary for coronavirus replication, because it seemed unlikely that HF could target a viral protein with similar efficacy for Alpha- and Betacoronavirus. A possible cellular target, HO-1, was identified. A recent study has shown that HO-1 pathway is activated upon treatment of melanoma cells with HF (39). Moreover, agonists of HO-1 pathway, such as hemin, are known to exhibit antiviral activity against SARS-CoV-2 (40). However, our results suggested that the antiviral activity of HF is not linked to HO-1 pathway. First, upregulation of HO-1 protein expression was not observed neither in Huh-7 cells nor in Vero81 cells upon HF treatment. It was shown by others that HO-1 expression could be induced in these cell types when treated with agonists such as hemin (40, 53–56). Second, the knock-down of HO-1 expression in Huh-7 cells did not impair HCoV-229E infection. Further investigations are needed to fully characterize the HF mechanism of action. Transcriptomic and proteomic analyses may help to identify cellular genes or proteins whose expression is regulated by HF. In conclusion, SJW extract and HF might be of great interest for future therapies on HCoV or animal coronaviruses. A proof of their efficacy *in vivo* is still needed. However, efficacy of HF against 4 different HCoVs makes this molecule particularly interesting in a pandemic preparedness approach, in the event of the emergence of a new highly pathogenic HCoV. Lately, new SARS-Like CoVs have been described in bats and are still a threat for human health (57, 58).

## MATERIALS AND METHODS

### Chemicals

Dulbecco’s modified Eagle’s medium (DMEM), phosphate-buffered saline (PBS), 4’,6-diamidino-2-phenylindole (DAPI), GlutaMAX^TM^ and Lipofectamine RNAi MAX were purchased from Life Technologies (Carlsbad, California, USA). Goat and fetal bovine sera (FBS) were obtained from Eurobio (Evry, France). Mowiol 4-88 was obtained from Calbiochem (Darmstadt, Germany). Remdesivir (GS-5734) and tariquidar were from BioTechne (Minneapolis, USA). Nirmatrelvir was from MedChemExpress (Monmouth Junction, USA). GC-376 was obtained from AmBeed (Arlington Heights, USA). Chloroquine was from Sigma-Aldrich (Saint Louis, USA). HC and HF were purchased from Phytolab (Vestenbergsgreuth, Germany) (total of HF > 98.0%). Pba was from Cayman Chemicals (Merck Chemicals, Darmstadt, Germany). Stocks of compounds were resuspended in dimethyl sulfoxide (DMSO) at 100 mM. Plant extracts were resuspended in DMSO at 50 mg/ml.

### Antibodies

Polyclonal rabbit anti-SARS-CoV-2 nucleocapsid antibody was purchased from Novus Biological (Cambridge, UK). Mouse anti-dsRNA mAb (clone J2) was from Scicons (Nordic-MUbio, Susteren, the Netherlands). Rabbit polyclonal anti-MERS-CoV Spike protein antibody was from SinoBiological (Eschborn, Germany). Mouse anti-ß-tubulin IgG1 antibody (T5201) was from Sigma. Monoclonal rabbit anti-HO-1 antibody was purchased from Cell Signaling Technology (Danvers, Massachusetts). Horseradish peroxidase-conjugated goat anti-rabbit IgG antibody, goat anti-mouse IgG antibody, Alexa 594-conjugated goat anti-rabbit antibody and Alexa 488-conjugated donkey anti-mouse antibody, were purchased from Jackson ImmunoResearch (Ely, United Kingdom).

### Cells

Human kidney cell lines (HEK293T/17, ATCC, CRL-11268; HEK293TT/ACE2) (59), African green monkey kidney cell lines (Vero-81, ATCC, CCL-81; Vero-E6 cells), Human lung cell line (A549/ACE2, kindly provided by Delphine Muriaux), and Human hepatoma cells (Huh-7) were grown in DMEM supplemented with 10% FBS.

Human lung cell line Calu-3 (ATCC, HTB-55) was cultivated in MEM supplemented with 10% FBS and glutaMAX-1.

Lentiviral vectors expressing TMPRSS2 were used to transduce Vero-81 cells and to produce Huh-7/TMPRSS2 stable cell line. This latter was selected with 2 µg/mL of puromycin. A reporter cell line, F1G-Red, generated in the laboratory, and derivates from Vero-81 cells was also used for combination assay (38). Primary human nasal epithelia MucilAir^TM^ (Epithelix, Geneva, Switzerland) were maintained in MucilAir^TM^ culture medium (Epithelix) as recommended by the manufacturer.

### Plant collection and extraction

The aerial part of *Hypericum perforatum* L. was collected in the Lille metropolis (France), deposited and identified at the Herbarium of the University of Lille by Gabriel Lefèvre and Céline Rivière. The plant was dried at 37°C for 48 to 72 h. After being powdered, the plant material was extracted three times with methanol for 24 h under agitation. The combined methanolic extract was then concentrated under reduced pressure with a rotary evaporator (Heidolph^TM^, Grosseron, Germany). The obtained crude methanolic extract was then resuspended with DMSO at the concentration of 50 mg/mL for the experiments.

### Virus

HCoV-229E-luc was kindly gifted by Volker Thiel (60). SARS-CoV-2 variants (the original Wuhan strain (EPI_ISL_410720), the alpha (B1.1.7; EPI_ISL_1653931) and omicron variants (B1.1.529; EPI_ISL_7696645). The original strain was kindly provided by the French National Reference Center for Respiratory Viruses hosted by Institut Pasteur (Paris, France). SARS-CoV strain (Frankfurt isolate) was provided by Dr Michelle Vialette (Unité de Sécurité Microbiologique). MERS-CoV was recovered by transfecting the infectious clone of MERS-CoV-EMC12 (kindly provided by Luis Enjuanes) in Huh-7 cells.

### Cell viability

Huh-7 or Vero-81 cells were seeded in 96-well plate at a density of 1x10^4^ and 1.5x10^4^ cells per well respectively, and incubated for 24 h at 37°C and 5% CO_2_. A 3-(4,5-dimethylthiazol-2-yl)-5-(3-carboxymethoxyphenyl)-2-(4-sulfophenyl)-2H-tetrazolium (MTS)-based viability assay (CellTiter 96® AQueous One Solution Cell Proliferation Assay, Promega) was performed as previously described (25).

### MucilAir^TM^ cytoxicity assays

Cytotoxicity was studied according to the manufacturer’s instruction either using cytotoxicity LDH assay kit-WST (Dojindo) or by measuring transepithelial electrical resistance (TEER) (Millicell® ERS-2, Millipore) as previously described (25). Toxicity is considered when LDH secretion is above 5% and TEER below 100 Ω.cm^2^.

### HCoVs infection assays

#### HCoV-229E-Luc

2x10^4^ Huh-7/TMPRSS2 cells per well were seeded into a 96-well plate 24 h before infection. Cells were inoculated with HCoV-229E-Luc and, simultaneously, increased concentrations of compound or the plant extract were added to cell culture medium. The inoculum was removed after 1 h and replaced with culture medium containing the compound or the plant extract. The cells were then lysed 7 h later in 20 µl of Renilla luciferase assay lysis buffer (Promega), and luciferase activity was quantified using a Tristar LB 941 luminometer (Berthold Technologies, Bad Bildbad, Germany) as recommended by the manufacturer.

#### MERS-CoV and SARS-CoV

2x10^5^ Calu-3 or Vero-81/TMPRSS2 cells were seeded in a 24-well plate on coverslips, 48 h or 24 h prior infection with MERS-CoV or SARS-CoV, respectively. Cells were inoculated with the virus at a multiplicity of infection (MOI) of 0.1, in the presence of increased concentrations of the compound, for 1 h at 37°C and 5% of CO_2_. The inoculum was replaced by culture medium containing the compound and the cells were incubated for 16 h. Supernatants were collected for viral titration and cells were fixed twice with 4% of paraformaldehyde (PFA) before exiting the BSL-3 facility and processed for immunostaining.

#### SARS-CoV-2

1x10^5^ Vero-81/TMPRSS2 or A549/ACE2 cells per well were seeded in a 48-well plate 24 h before infection. Cells were inoculated with the virus at a MOI of 0.3, in the presence of increased concentration of the compound, for 1 h at 37°C and 5% of CO_2_. 50 nM tariquidar, a P-glycoprotein inhibitor, was added in the media to inhibit pump efflux, and 10 µM chloroquine was used as a control of the expression of TMPRSS2. Inoculum was replaced with media containing the different compounds and cells were incubated for 16 h at 37°C and 5% of CO_2_. The supernatants were collected for viral titration and the cells were lysed using reducing Laëmmli loading buffer for western blot analysis. The samples were inactivated 30 min at 95°C.

### Light-dependent assay

Cells were inoculated with HCoV-229E-Luc, treated with increasing concentrations of the compound and incubated in the cabinet under light or dark conditions before being place in the incubator at 37°C and 5% CO_2_. For the light condition, the plate was incubated for 10 min under the light of the safety cabinet. For the dark condition, the experiment was conducted in the dark, with the light of the safety cabinet turned off, and the tubes and plate were covered with aluminum foil. Only the light from the hood next to the safety cabinet was on. After 7 h of incubation, cells were lysed and infection was studied by measuring the luciferase activity.

### Infectivity titration

Huh-7 (MERS-CoV) or Vero-E6 (SARS-CoV and SARS-CoV-2) were seeded in a 96-well plate and were inoculated with 1/10 serially diluted supernatants. After 5 days (SARS-CoV and SARS-CoV-2) or 7 days (MERS-CoV) of incubation at 37°C and 5% of CO_2_, the 50% tissue culture infectious dose (TCID50/mL) was determined by assessing the virus-induced cytopathic effect and using the Spearman-Kärber formula.

### Western blot detection

Proteins were separated onto a 12% SDS-polyacrylamide gel electrophoresis and transferred on nitrocellulose membranes (Hybond-ECL, Amersham). The membranes were blocked and incubated overnight at 4°C with a polyclonal rabbit anti-SARS-CoV-2 nucleocapsid antibody (1/4000), a mouse anti-ß-tubulin (1/4000) or a rabbit anti-HO-1 (1/1000). They were visualized by enhanced chemoluminescence (Pierce^TM^ ECL, ThermoFisher Scientific) on LAS3000 (Fujifilm) or Amersham ImageQuant 800 (Cytiva).

### Immunostaining

Cells were permeabilized for 5 min with 0.4 % Triton X-100 and blocked with 5% of goat serum for 30 min and were incubated with anti-dsRNA monoclonal mouse antibody (clone J2) or anti-MERS-CoV Spike protein polyclonal rabbit antibody. Cells were rinsed three times with PBS, and immunostained with an Alexa 594-conjugated goat anti-rabbit secondary antibody or an alexa-488-conjugated donkey anti-mouse secondary antibody and DAPI. The coverslips were mounted on microscope slides in Mowiol. The images were acquired with an Evos M5000 microscope (Thermo Fischer Scientific). Ten images were randomly taken for each condition in duplicate. The number of cells were determined by the number of nuclei, and infected cells were detected by quantifying the number of dsRNA-positive or Spike-positive cells.

### MucilAir^TM^-Human airway epithelia (MucilAir^TM^-HAE) infection assay

The apical surface of the cells was rinsed 3 times for 10 min using MucilAir^TM^-HAE culture medium to remove the mucosal secretion. The cells were inoculated at the apical side with HCoV-229E-Luc (MOI = 0.01) or SARS-CoV-2 (MOI = 0.3) and treated with 4 µM or 12 µM or HF or 0.025% DMSO for 1 h. On the apical pole, the inoculum was removed and replaced by 10 µL of medium containing the compounds. Simultaneously, HF or DMSO were added in the basolateral medium.

For HCoV-229E-Luc infection, after 24 h of incubation, 140 µL of culture medium was added on the apical surface of MucilAir^TM^-HAE and collected for RNA extraction. The cells were then lysed with 40 µL of Renilla luciferase assay lysis buffer (Promega). Luciferase activity was quantified as previously described.

For SARS-CoV-2 infection, after 48 h of incubation, 140 µL of culture medium was added on the apical surface of MucilAir^TM^-HAE and was collected for RNA extraction and viral titration. The cells were lysed and RNA was extracted for RT-qPCR assay.

### RT-qPCR assay

RNA was extracted from MucilAir^TM^-HAE supernatants or cells using QIAamp Viral RNA Mini kit (Qiagen) and NucleoSpin RNA plus (Macherey Nagel) respectively. One-step qPCR assay was performed using 5 µL of RNA and Takyon Low rox one-step RT probe master mix (UFD-LPRT-C0101, Eurogentec) with specific primers and probes (**Supplemental Table 1**) and using a Quantstudio 3 (Applied Biosystems). The expressions of HCoV-229E M gene and SARS-CoV-2 E gene were quantified using a standard curve.

### Time-of-addition assay

One day prior the infection, 1x10^5^ Vero-81/TMPRSS2 cells per well and 2x10^4^ Huh-7/TMPRSS2 cells per well were seeded into a 48-well plate or 96-well plate for SARS-CoV-2 or HCoV-229E-Luc infection, respectively. To assess which viral step is inhibited, the different compounds were added at different time points, either 1 h before the inoculation (corresponding to the condition “pre-treatment”), or during the inoculation or 1 h, 2 h, 3 h after the inoculation (1 h p.i., 2 h p.i. or 3 h p.i.). The cells were then lysed at 16 h or 7 h post-infection for SARS-CoV-2 or HCoV-229E-Luc, respectively, and analyzed as described ahead.

### Pseudotyped particle entry assay

Particles pseudotyped with either the spike protein of SARS-CoV-2 (SARS2pp), or HCoV-229E (HCoV-229Epp), were produced as previously described (61). 4.5x10^3^ HEK293TT/ACE2 (SARS2pp) cells per well or 1x10^4^ Huh-7/TMPRSS2 (229Epp) cells per well were seeded into a 96-well plate 24 h before infection. Cells were inoculated with SARS2pp and 229Epp in the presence of HF or HC (2.5, 5 or 10 µM) for 2 h at 37°C and 5% of CO_2_. The inoculum was removed and replaced with culture media without compound. The cells were lysed after 48 h of incubation. The luciferase activity was then measured with Firefly luciferase assay kit (Promega) according to the manufacturer recommendations and using a luminometer (Berthold).

### Drug combination assay

24 h before infection, 4.5x10_3_ F1G-Red cells per well were seeded in 384-well plate in DMEM with 2% of FBS. Compounds (HF, remdesivir and nirmatrelvir) were dissolved in DMSO at concentrations stocks of 10 or 100 nM. 1 h before infection, dose-response concentrations of the compound (seven-2-fold serial dilutions) were dispensed onto the cells using an Echo 550 acoustic dispenser (Labcyte) in three biological replicates. Last column of each plate contained DMSO control solvent. DMSO was distributed at equivalent volumes as negative control. The cells were then inoculated with the virus by adding 10 µL of inoculum (MOI = 0.3) and 50 nM tariquidar for 16 h. The infection was assessed by using an IN CELL Analyzer 6500 high-throughput automated confocal microscope (Ge Healthcare) located in BSL-3 safety laboratory. Nine images were taken (20X objective, NA 0.75) for each condition and the number of infected cells was quantified using Columbus image analysis software (Perkin Elmer). The data were analyzed and we used Synergy Finder (http://www.synergyfinderplus.org/) for the calculation of the synergy scores (mean and *p*-value) and https://synergyfinder.fimm.fi/ for the most synergistic area (MSA).

### RNA interference

Cells were transfected with small interfering RNA (siRNA) targeting HO-1 (siHO-1, final concentration: 10 nM) (Gene HMOX1, AM16708, Assay ID 11056, ThermoFischer) or with a non-targeting siRNA control (siCTRL, Dharmacon) using Lipofectamine RNAi MAX (Invitrogen) according to manufacturer’s recommendations. After 24 h of incubation, cells were treated with increasing concentration of hemin or HF, infected or not with HCoV-229E-Luc and incubated again for 24 h. Cells were then lysed using Laëmmli buffer in reducing conditions or with the Renilla luciferase assay lysis buffer. HO-1 expression was then studied by Western Blot.

### Statistical analysis and IC_50_ and CC_50_ calculation

IC_50_ and CC_50_ values were calculated by nonlinear regression curve fitting with variable slopes and mean and standard error of the mean (SEM) values were graphed using GraphPad Prism software version 10.0.3 (Boston, Ma, USA) and by constraining the top to 100% and the bottom to 0%. Statistical analysis was performed using Mann-Whitney non-parametric test by comparing each treated group with the untreated control (DMSO control). *P*-values < 0.05 were considered significantly different from the control.

## ACKNOWLEDGMENTS

We are grateful for the technical help of Robin Prath and Nicolas Vandenabeele in the BSL-3 facility. We also thank the team of Fernando Real, especially Cyrine Bentaleb for useful discussions. We also would like to thank Steve Polyak for his advices on combination assays. Finally, we would like to thank Yves Rouillé for kindly supplying F1G-Red cell line. We thank Lola Dandoy for technical help.

This project was funded by Région Hauts-de-France and I-Site (FlavoCoV project) and CNRS (VIROCRIB project). We thank the Fédération Hospitalo-Universitaire (FHU) RESPIRE for financial support.

I.R. is a recipient of a Health-PhD fellowship. L.D. is a recipient of a CNRS and Institut Pasteur de Lille fellowship.

## REFERENCES

1. Owen DR, Allerton CMN, Anderson AS, Aschenbrenner L, Avery M, Berritt S, Boras B, Cardin RD, Carlo A, Coffman KJ, Dantonio A, Di L, Eng H, Ferre R, Gajiwala KS, Gibson SA, Greasley SE, Hurst BL, Kadar EP, Kalgutkar AS, Lee JC, Lee J, Liu W, Mason SW, Noell S, Novak JJ, Obach RS, Ogilvie K, Patel NC, Pettersson M, Rai DK, Reese MR, Sammons MF, Sathish JG, Singh RSP, Steppan CM, Stewart AE, Tuttle JB, Updyke L, Verhoest PR, Wei L, Yang Q, Zhu Y. 2021. An oral SARS-CoV-2 Mpro inhibitor clinical candidate for the treatment of COVID-19. Science 374:1586–1593.

2. Lamb YN. 2022. Nirmatrelvir Plus Ritonavir: First Approval. Drugs 82:585–591.

3. Warren TK, Jordan R, Lo MK, Ray AS, Mackman RL, Soloveva V, Siegel D, Perron M, Bannister R, Hui HC, Larson N, Strickley R, Wells J, Stuthman KS, Van Tongeren SA, Garza NL, Donnelly G, Shurtleff AC, Retterer CJ, Gharaibeh D, Zamani R, Kenny T, Eaton BP, Grimes E, Welch LS, Gomba L, Wilhelmsen CL, Nichols DK, Nuss JE, Nagle ER, Kugelman JR, Palacios G, Doerffler E, Neville S, Carra E, Clarke MO, Zhang L, Lew W, Ross B, Wang Q, Chun K, Wolfe L, Babusis D, Park Y, Stray KM, Trancheva I, Feng JY, Barauskas O, Xu Y, Wong P, Braun MR, Flint M, McMullan LK, Chen S-S, Fearns R, Swaminathan S, Mayers DL, Spiropoulou CF, Lee WA, Nichol ST, Cihlar T, Bavari S. 2016. Therapeutic efficacy of the small molecule GS-5734 against Ebola virus in rhesus monkeys. Nature 531:381–385.

4. Agostini ML, Andres EL, Sims AC, Graham RL, Sheahan TP, Lu X, Smith EC, Case JB, Feng JY, Jordan R, Ray AS, Cihlar T, Siegel D, Mackman RL, Clarke MO, Baric RS, Denison MR. 2018. Coronavirus Susceptibility to the Antiviral Remdesivir (GS-5734) Is Mediated by the Viral Polymerase and the Proofreading Exoribonuclease. mBio 9:10.1128/mbio.00221-18.

5. Beigel JH, Tomashek KM, Dodd LE, Mehta AK, Zingman BS, Kalil AC, Hohmann E, Chu HY, Luetkemeyer A, Kline S, Lopez de Castilla D, Finberg RW, Dierberg K, Tapson V, Hsieh L, Patterson TF, Paredes R, Sweeney DA, Short WR, Touloumi G, Lye DC, Ohmagari N, Oh M-D, Ruiz-Palacios GM, Benfield T, Fätkenheuer G, Kortepeter MG, Atmar RL, Creech CB, Lundgren J, Babiker AG, Pett S, Neaton JD, Burgess TH, Bonnett T, Green M, Makowski M, Osinusi A, Nayak S, Lane HC, ACTT-1 Study Group Members. 2020. Remdesivir for the Treatment of Covid-19 - Final Report. N Engl J Med 383:1813– 1826.

6. Yin W, Mao C, Luan X, Shen D-D, Shen Q, Su H, Wang X, Zhou F, Zhao W, Gao M, Chang S, Xie Y-C, Tian G, Jiang H-W, Tao S-C, Shen J, Jiang Y, Jiang H, Xu Y, Zhang S, Zhang Y, Xu HE. 2020. Structural basis for inhibition of the RNA-dependent RNA polymerase from SARS-CoV-2 by remdesivir. Science 368:1499–1504.

7. Sheahan TP, Sims AC, Zhou S, Graham RL, Pruijssers AJ, Agostini ML, Leist SR, Schäfer A, Dinnon KH, Stevens LJ, Chappell JD, Lu X, Hughes TM, George AS, Hill CS, Montgomery SA, Brown AJ, Bluemling GR, Natchus MG, Saindane M, Kolykhalov AA, Painter G, Harcourt J, Tamin A, Thornburg NJ, Swanstrom R, Denison MR, Baric RS. 2020. An orally bioavailable broad-spectrum antiviral inhibits SARS-CoV-2 in human airway epithelial cell cultures and multiple coronaviruses in mice. Science Translational Medicine 12:eabb5883.

8. Stuyver LJ, Whitaker T, McBrayer TR, Hernandez-Santiago BI, Lostia S, Tharnish PM, Ramesh M, Chu CK, Jordan R, Shi J, Rachakonda S, Watanabe KA, Otto MJ, Schinazi RF. 2003. Ribonucleoside Analogue That Blocks Replication of Bovine Viral Diarrhea and Hepatitis C Viruses in Culture. Antimicrobial Agents and Chemotherapy 47:244–254.

9. Uraki R, Ito M, Kiso M, Yamayoshi S, Iwatsuki-Horimoto K, Furusawa Y, Sakai-Tagawa Y, Imai M, Koga M, Yamamoto S, Adachi E, Saito M, Tsutsumi T, Otani A, Kikuchi T, Yotsuyanagi H, Halfmann PJ, Pekosz A, Kawaoka Y. 2023. Antiviral and bivalent vaccine efficacy against an omicron XBB.1.5 isolate. Lancet Infect Dis 23:402–403.

10. Imai M, Ito M, Kiso M, Yamayoshi S, Uraki R, Fukushi S, Watanabe S, Suzuki T, Maeda K, Sakai-Tagawa Y, Iwatsuki-Horimoto K, Halfmann PJ, Kawaoka Y. 2023. Efficacy of Antiviral Agents against Omicron Subvariants BQ.1.1 and XBB. N Engl J Med 388:89– 91.

11. Bez P, D’ippolito G, Deiana CM, Finco Gambier R, Pica A, Costanzo G, Garzi G, Scarpa R, Landini N, Cinetto F, Firinu D, Milito C. 2023. Struggling with COVID-19 in Adult Inborn Errors of Immunity Patients: A Case Series of Combination Therapy and Multiple Lines of Therapy for Selected Patients. 7. Life 13:1530.

12. Baldi F, Dentone C, Mikulska M, Fenoglio D, Mirabella M, Magnè F, Portunato F, Altosole T, Sepulcri C, Giacobbe DR, Uras C, Scavone G, Taramasso L, Orsi A, Cittadini G, Filaci G, Bassetti M. 2023. Case report: Sotrovimab, remdesivir and nirmatrelvir/ritonavir combination as salvage treatment option in two immunocompromised patients hospitalized for COVID-19. Frontiers in Medicine 9.

13. Brown L-AK, Moran E, Goodman A, Baxendale H, Bermingham W, Buckland M, AbdulKhaliq I, Jarvis H, Hunter M, Karanam S, Patel A, Jenkins M, Robbins A, Khan S, Simpson T, Jolles S, Underwood J, Savic S, Richter A, Shields A, Brown M, Lowe DM. 2022. Treatment of chronic or relapsing COVID-19 in immunodeficiency. Journal of Allergy and Clinical Immunology 149:557–561.e1.

14. Anderson AS, Caubel P, Rusnak JM, EPIC-HR Trial Investigators. 2022. Nirmatrelvir-Ritonavir and Viral Load Rebound in Covid-19. N Engl J Med 387:1047–1049.

15. Charness ME, Gupta K, Stack G, Strymish J, Adams E, Lindy DC, Mohri H, Ho DD. 2022. Rebound of SARS-CoV-2 Infection after Nirmatrelvir–Ritonavir Treatment. N Engl J Med 387:1045–1047.

16. EMA. 2023. Lagevrio: Withdrawn application. European Medicines Agency. Text. https://www.ema.europa.eu/en/medicines/human/withdrawn-applications/lagevrio. Retrieved 22 September 2023.

17. Zhou S, Hill CS, Sarkar S, Tse LV, Woodburn BMD, Schinazi RF, Sheahan TP, Baric RS, Heise MT, Swanstrom R. 2021. β-d-N4-hydroxycytidine Inhibits SARS-CoV-2 Through Lethal Mutagenesis But Is Also Mutagenic To Mammalian Cells. J Infect Dis 224:415– 419.

18. Coronaviridae Study Group of the International Committee on Taxonomy of Viruses. 2020. The species Severe acute respiratory syndrome-related coronavirus: classifying 2019-nCoV and naming it SARS-CoV-2. Nat Microbiol 5:536–544.

19. Tang G, Liu Z, Chen D. 2022. Human coronaviruses: Origin, host and receptor. Journal of Clinical Virology 155:105246.

20. Belouzard S, Millet JK, Licitra BN, Whittaker GR. 2012. Mechanisms of Coronavirus Cell Entry Mediated by the Viral Spike Protein. Viruses 4:1011–1033.

21. Jackson CB, Farzan M, Chen B, Choe H. 2022. Mechanisms of SARS-CoV-2 entry into cells. Nat Rev Mol Cell Biol 23:3–20.

22. Hoffmann M, Pöhlmann S. 2021. How SARS-CoV-2 makes the cut. Nat Microbiol 6:828– 829.

23. Weiss SR, Leibowitz JL. 2011. Coronavirus Pathogenesis. Adv Virus Res 81:85–164.

24. Denaro M, Smeriglio A, Barreca D, De Francesco C, Occhiuto C, Milano G, Trombetta D. 2020. Antiviral activity of plants and their isolated bioactive compounds: An update. Phytother Res 34:742–768.

25. Meunier T, Desmarets L, Bordage S, Bamba M, Hervouet K, Rouillé Y, François N, Decossas M, Sencio V, Trottein F, Tra Bi FH, Lambert O, Dubuisson J, Belouzard S, Sahpaz S, Séron K. 2022. A Photoactivable Natural Product with Broad Antiviral Activity against Enveloped Viruses, Including Highly Pathogenic Coronaviruses. Antimicrob Agents Chemother 66:e0158121.

26. Millet JK, Séron K, Labitt RN, Danneels A, Palmer KE, Whittaker GR, Dubuisson J, Belouzard S. 2016. Middle East respiratory syndrome coronavirus infection is inhibited by griffithsin. Antiviral Res 133:1–8.

27. Al Ibrahim M, Akissi ZLE, Desmarets L, Lefèvre G, Samaillie J, Raczkiewicz I, Sahpaz S, Dubuisson J, Belouzard S, Rivière C, Séron K. 2023. Discovery of Anti-Coronavirus Cinnamoyl Triterpenoids Isolated from Hippophae rhamnoides during a Screening of Halophytes from the North Sea and Channel Coasts in Northern France. Int J Mol Sci 24:16617.

28. Hypericum Tourn. ex L. | Plants of the World Online | Kew Science. Plants of the World Online. http://powo.science.kew.org/taxon/urn:lsid:ipni.org:names:30002180-2. Retrieved 18 November 2022.

29. de Carvalho Meirelles G, Bridi H, von Poser GL, Nemitz MC. 2019. Hypericum species: An analysis on the patent technologies. Fitoterapia 139:104363.

30. Butterweck V, Schmidt M. 2007. St. John’s wort: role of active compounds for its mechanism of action and efficacy. Wien Med Wochenschr 157:356–361.

31. EMA. 2023. Hyperici herba. European Medicines Agency. Text. https://www.ema.europa.eu/en/medicines/herbal/hyperici-herba-0. Retrieved 9 April 2024.

32. Röder C, Schaefer M, Leucht S. 2004. [Meta-analysis of effectiveness and tolerability of treatment of mild to moderate depression with St. John’s Wort]. Fortschr Neurol Psychiatr 72:330–343.

33. Linde K, Berner MM, Kriston L. 2008. St John’s wort for major depression. Cochrane Database Syst Rev 2008:CD000448.

34. Kubin A, Wierrani F, Burner U, Alth G, Grünberger W. 2005. Hypericin--the facts about a controversial agent. Curr Pharm Des 11:233–253.

35. Lopez-Bazzocchi I, Hudson JB, Towers GHN. 1991. Antiviral Activity of the Photoactive Plant Pigment Hypericin. Photochemistry and Photobiology 54:95–98.

36. Mohamed FF, Anhlan D, Schöfbänker M, Schreiber A, Classen N, Hensel A, Hempel G, Scholz W, Kühn J, Hrincius ER, Ludwig S. 2022. Hypericum perforatum and Its Ingredients Hypericin and Pseudohypericin Demonstrate an Antiviral Activity against SARS-CoV-2. Pharmaceuticals (Basel) 15:530.

37. Jendželovská Z, Jendželovský R, Kuchárová B, Fedoročko P. 2016. Hypericin in the Light and in the Dark: Two Sides of the Same Coin. Front Plant Sci 7:560.

38. Desmarets L, Callens N, Hoffmann E, Danneels A, Lavie M, Couturier C, Dubuisson J, Belouzard S, Rouillé Y. 2022. A reporter cell line for the automated quantification of SARS-CoV-2 infection in living cells. Front Microbiol 13:1031204.

39. Cardile A, Passarini C, Zanrè V, Fiore A, Menegazzi M. 2023. Hyperforin Enhances Heme Oxygenase-1 Expression Triggering Lipid Peroxidation in BRAF-Mutated Melanoma Cells and Hampers the Expression of Pro-Metastatic Markers. Antioxidants (Basel) 12:1369.

40. Kim D-H, Ahn H-S, Go H-J, Kim D-Y, Kim J-H, Lee J-B, Park S-Y, Song C-S, Lee S-W, Ha S-D, Choi C, Choi I-S. 2021. Hemin as a novel candidate for treating COVID-19 via heme oxygenase-1 induction. Sci Rep 11:21462.

41. Espinoza JA, González PA, Kalergis AM. 2017. Modulation of Antiviral Immunity by Heme Oxygenase-1. Am J Pathol 187:487–493.

42. Maridass M, Britto AD. 2008. Origins of Plant Derived Medicines. Ethnobotanical Leaflets 2008.

43. Wang L, Zhang R-M, Liu G-Y, Wei B-L, Wang Y, Cai H-Y, Li F-S, Xu Y-L, Zheng S-P, Wang G. 2010. Chinese herbs in treatment of influenza: A randomized, double-blind, placebo-controlled trial. Respiratory Medicine 104:1362–1369.

44. Adel Mehraban MS, Shirzad M, Mohammad Taghizadeh Kashani L, Ahmadian-Attari MM, Safari AA, Ansari N, Hatami H, Kamalinejad M. 2023. Efficacy and safety of add-on Viola odorata L. in the treatment of COVID-19: A randomized double-blind controlled trial. J Ethnopharmacol 304:116058.

45. Wilson D, Goggin K, Williams K, Gerkovich MM, Gqaleni N, Syce J, Bartman P, Johnson Q, Folk WR. 2015. Consumption of Sutherlandia frutescens by HIV-Seropositive South African Adults: An Adaptive Double-Blind Randomized Placebo Controlled Trial. PLoS One 10:e0128522.

46. Hudson JB, Lopez-Bazzocchi I, Towers GH. 1991. Antiviral activities of hypericin. Antiviral Res 15:101–112.

47. Barnes J, Anderson LA, Phillipson JD. 2001. St John’s wort (Hypericum perforatum L.): a review of its chemistry, pharmacology and clinical properties. J Pharm Pharmacol 53:583–600.

48. Biber A, Fischer H, Römer A, Chatterjee SS. 1998. Oral bioavailability of hyperforin from hypericum extracts in rats and human volunteers. Pharmacopsychiatry 31 Suppl 1:36–43.

49. Hatanaka J, Shinme Y, Kuriyama K, Uchida A, Kou K, Uchida S, Yamada S, Onoue S. 2011. In vitro and in vivo characterization of new formulations of St. John’s Wort extract with improved pharmacokinetics and anti-nociceptive effect. Drug Metab Pharmacokinet 26:551–558.

50. Agrosí M, Mischiatti S, Harrasser PC, Savio D. 2000. Oral bioavailability of active principles from herbal products in humans. A study on *Hypericum perforatum* extracts using the soft gelatin capsule technology. Phytomedicine 7:455–462.

51. Negreş S, Scutari C, Ionică FE, Gonciar V, Velescu BŞ, Şeremet OC, Zanfirescu A, Zbârcea CE, Ştefănescu E, Ciobotaru E, ChiriŢă C. 2016. Influence of hyperforin on the morphology of internal organs and biochemical parameters, in experimental model in mice. Rom J Morphol Embryol 57:663–673.

52. Donà M, Dell’Aica I, Pezzato E, Sartor L, Calabrese F, Barbera MD, Donella-Deana A, Appendino G, Borsarini A, Caniato R, Garbisa S. 2004. Hyperforin Inhibits Cancer Invasion and Metastasis. Cancer Research 64:6225–6232.

53. Hill-Batorski L, Halfmann P, Neumann G, Kawaoka Y. 2013. The cytoprotective enzyme heme oxygenase-1 suppresses Ebola virus replication. J Virol 87:13795–13802.

54. Zhu Z, Wilson AT, Mathis MM, Wen F, Brown KE, Luxon BA, Schmidt WN. 2008. Heme Oxygenase-1 suppresses Hepatitis C Virus replication and increases resistance of hepatocytes to oxidant injury. Hepatology 48:1430–1439.

55. Tseng C-K, Lin C-K, Wu Y-H, Chen Y-H, Chen W-C, Young K-C, Lee J-C. 2016. Human heme oxygenase 1 is a potential host cell factor against dengue virus replication. 1. Sci Rep 6:32176.

56. Ibáñez FJ, Farías MA, Retamal-Díaz A, Espinoza JA, Kalergis AM, González PA. 2017. Pharmacological Induction of Heme Oxygenase-1 Impairs Nuclear Accumulation of Herpes Simplex Virus Capsids upon Infection. Front Microbiol 8:2108.

57. Li W, Shi Z, Yu M, Ren W, Smith C, Epstein JH, Wang H, Crameri G, Hu Z, Zhang H, Zhang J, McEachern J, Field H, Daszak P, Eaton BT, Zhang S, Wang L-F. 2005. Bats are natural reservoirs of SARS-like coronaviruses. Science 310:676–679.

58. Temmam S, Vongphayloth K, Baquero E, Munier S, Bonomi M, Regnault B, Douangboubpha B, Karami Y, Chrétien D, Sanamxay D, Xayaphet V, Paphaphanh P, Lacoste V, Somlor S, Lakeomany K, Phommavanh N, Pérot P, Dehan O, Amara F, Donati F, Bigot T, Nilges M, Rey FA, van der Werf S, Brey PT, Eloit M. 2022. Bat coronaviruses related to SARS-CoV-2 and infectious for human cells. Nature 604:330–336.

59. Lavie M, Dubuisson J, Belouzard S. 2022. SARS-CoV-2 Spike Furin Cleavage Site and S2’ Basic Residues Modulate the Entry Process in a Host Cell-Dependent Manner. J Virol 96:e0047422.

60. van den Worm SHE, Eriksson KK, Zevenhoven JC, Weber F, Züst R, Kuri T, Dijkman R, Chang G, Siddell SG, Snijder EJ, Thiel V, Davidson AD. 2012. Reverse genetics of SARS-related coronavirus using vaccinia virus-based recombination. PLoS One 7:e32857.

61. Belouzard S, Chu VC, Whittaker GR. 2009. Activation of the SARS coronavirus spike protein via sequential proteolytic cleavage at two distinct sites. Proc Natl Acad Sci U S A 106:5871–5876.

